# Leveraging Single-Cell Sequencing to Classify and Characterize Tumor Subgroups in Bulk RNA-Sequencing Data

**DOI:** 10.1101/2024.03.02.583114

**Authors:** Arya Shetty, Su Wang, A. Basit Khan, Collin W. English, Shervin Hosseingholi Nouri, Stephen T. Magill, David R. Raleigh, Tiemo J. Klisch, Arif O. Harmanci, Akash J. Patel, Akdes Serin Harmanci

## Abstract

Accurate classification of cancer subgroups is essential for precision medicine, tailoring treatments to individual patients based on their cancer subtypes. In recent years, advances in high-throughput sequencing technologies have enabled the generation of large-scale transcriptomic data from cancer samples. These data have provided opportunities for developing computational methods that can improve cancer subtyping and enable better personalized treatment strategies. Here in this study, we evaluated different feature selection schemes in the context of meningioma classification. While the scheme relying solely on bulk transcriptomic data showed good classification accuracy, it exhibited confusion between malignant and benign molecular classes in approximately ~8% of meningioma samples. In contrast, models trained on features learned from meningioma single-cell data accurately resolved the sub-groups confused by bulk-transcriptomic data but showed limited overall accuracy. To integrate interpretable features from the bulk (n=78 samples) and single-cell profiling (~10K cells), we developed an algorithm named CLIPPR which combines the top-performing single-cell models with RNA-inferred copy number variation (CNV) signals and the initial bulk model to create a meta-model, which exhibited the strongest performance in meningioma classification. CLIPPR showed superior overall accuracy and resolved benign-malignant confusion as validated on n=792 bulk meningioma samples gathered from multiple institutions. Finally, we showed the generalizability of our algorithm using our in-house single-cell (~200K cells) and bulk TCGA glioma data (n=711 samples). Overall, our algorithm CLIPPR synergizes the resolution of single-cell data with the depth of bulk sequencing and enables improved cancer sub-group diagnoses and insights into their biology.

## Introduction

Brain tumors are highly heterogeneous neoplastic tissues^1^. Due to this complexity, the precision medicine-based treatment approaches rely on the classification of tumors into several categories that were shown to correlate with their prognosis, and treatment outcomes^2^. The WHO provides classifications and guidelines that are crucial for treatment decisions^3^. In many cases, molecular profiling, notably transcriptional and genetic data, outperforms histopathology-based classifications for brain tumors^4^. While numerous cancers are well-characterized, there is still a need to develop new methods that can help clinicians and researchers optimize molecular classification methods. Furthermore, recent years have brought about a surge of bulk and single-cell transcriptomic datasets. These rich datasets can further refine tumor classifications with the help of machine learning and statistical methods. Here, our goal is to develop a computational method for tumor classification, starting with meningioma tumor classification as a test case. This will enable a deep-dive analysis of the reasons underpinning computational misclassifications between different subgroups. We develop our pipeline by focusing on features extracted from bulk and single-cell transcriptomic datasets. Our focus on meningioma is validated by two key factors: Firstly, the availability of abundant in-house and publicly available datasets for this study. Secondly, meningioma is a well-characterized tumor that can be extensively studied in a computational deep-dive approach.

Meningiomas are the most common primary intracranial tumors^5^. Though typically benign, aggressive cases are not uncommon^6^. Classically, the WHO grading system has been used to classify these tumors and guide clinical management^3,7^. In our previous work, we employed an integrated multi-omics profiling approach to study the classification of human meningiomas^4^. Our analysis and others led to the identification of three distinct molecular groups: A, B, and C, in order of worsening prognoses^8–10^. We have confirmed the presence of these three molecular subgroups on various molecular profiling platforms, across species^11–13^ and demonstrated that they provide a more accurate prediction of the risk of recurrence when compared to the conventional World Health Organization (WHO) grading system comprising three grades (I, II, and III) ^8,9,14,15^. MenG A meningiomas are benign and show low levels of chromosomal instability. MenG B meningioma tend to have merlin loss, either via NF2 mutation or chr 22q loss^16–18^. MenG C tumors, the most aggressive, have NF2 mutations or chr22q loss and chromosomal instability, most commonly chr1p loss^9,19^. Given their clinically distinct presentations, which better predict long-term outcomes than the current WHO grading system, this classification scheme represents a promising new paradigm to guide future meningioma therapy^20^. Despite its promise, the full clinical potential of this classification scheme has yet to be fully realized.

While our molecular meningioma classifications are consistent and reproducible across publicly available datasets, classifiers that rely solely on bulk transcriptomic data have demonstrated confusion in approximately 8% of patient samples with benign (MenG A) and malignant (MenG C) tumors^9^. This confusion likely results from the high-dimensional nature of these classes, which makes it challenging to establish strictly delineated class definitions. Samples are characterized by examining the aggregated expression of hundreds of genes. As a result, the expression profile of samples within a given class can vary across a continuous spectrum. Furthermore, bulk-transcriptomics suffers from a loss of resolution that can obscure the molecular signature that would otherwise be used to assign a sample’s class^21^. The issues surrounding the accurate identification of clinically distinct groups stand in the way of implementing molecular classification schemes as a diagnostic tool in the management of meningioma. As such, optimizing molecular diagnostic tools stands as a high-priority target for further study.

Here we developed CLIPPR, a method that predicts the meningioma classes by leveraging single-cell data and RNA-inferred CNV signal to enhance the prediction accuracy of bulk data classifiers. We demonstrate that using models trained on features learned from single-cell data accurately resolved the confusion between the benign MenG A and the malignant MenG C groups but had limited overall accuracy. Similarly, models generated from RNA-inferred large-scale CNV signals also predicted malignant class accurately with limited overall accuracy. However, a meta-model-based combination of the top-performing single-cell models, CNV models, and the initial bulk model into a meta-model resulted in the strongest performance, with superior overall accuracy and benign-malignant resolution. We validated the algorithm’s generalizability by applying it to bulk TCGA RNA-Seq glioma data, incorporating features extracted from both the training set of bulk TCGA RNA-Seq glioma data and our in-house single-cell glioma data^22^.

## Results

### Overview of CLIPPR algorithm

The overview of the CLIPPR algorithm is shown in **Figure 1**. The input for training CLIPPR models consists of aligned single-cell and bulk RNA-seq read counts, as well as the bulk RNA-seq training cohort sample names and tumor classes (MenG groups). Additionally, training the model requires the tumor classes and cell types for each cell within the single-cell data. We assume that these classes are assigned by expert determination and by long-term follow-ups from an existing cohort.

**Figure 1.**
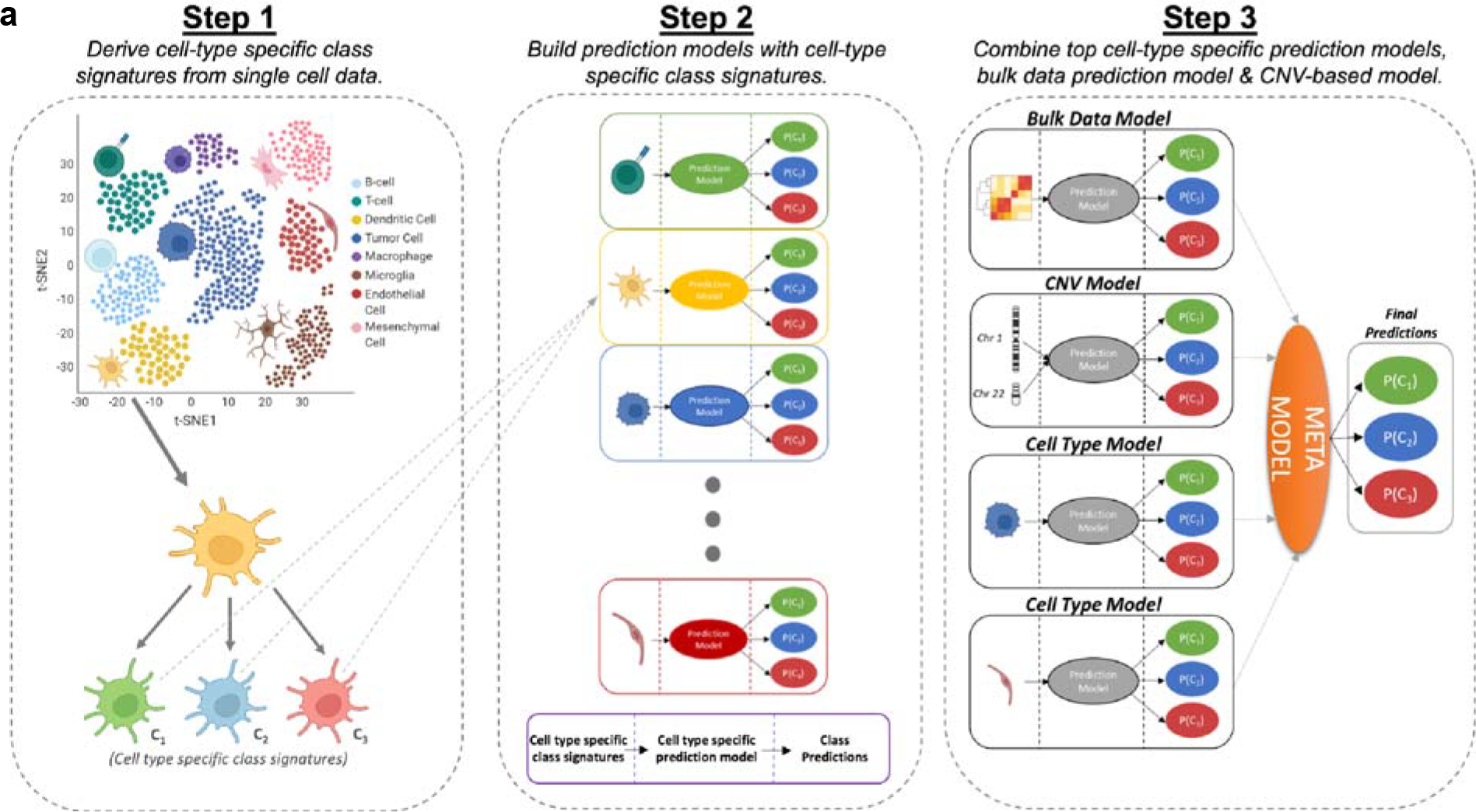
Overview of the CLIPPR algorithm. **(a)** Step 1 involves deriving cell type-specific class signatures from single-cell meningioma data. In Step 2, prediction models are constructed using cell type-specific class features (ctRFs). For each feature set, the bulk RNA-Seq data was used to train a cell-type specific Random Forest model. Models were also constructed using RNA-inferred CNV signals (cnvRF) and the bulk model (bulkRF). In Step 3, a meta-model is assembled, integrating the top cell type-specific models, the bulk model, and the CNV model.

The CLIPPR algorithm aims to leverage insights from both bulk transcriptomic and single-cell sequencing to generate high-performing models for the accurate classification of bulk transcriptomic sequencing. To this end, CLIPPR utilizes three distinct sets of features: a) Differentially expressed genes specific to each subtype derived from bulk RNA-Seq data. b) Differentially expressed genes that are specific to both cell type and subtype, extracted from single-cell RNA-Seq. c) CNV profiles of chromosomes of interest inferred from bulk transcriptomics.

#### Bulk RNA-Seq based model (bulkRF)

We trained a baseline bulk-transcriptomic model using the differentially expressed genes (DEGs) specific to each class within the well-characterized *bulk RNA-Seq* meningioma samples in the training cohort. These DEGs were used to train a Random Forest classifier.

#### ScRNA-Seq based cell type-specific models (ctRFs)

To leverage single-cell sequencing, we identified cell-type specific, class-specific differentially expressed genes (scDEGs). The bulk RNA-Seq sequencing corresponding to each set of scDEGs genes was then used to train cell-type specific Random Forest model (ctRFs).

#### CNV-Based Model (cnvRF)

MenG C, the most aggressive form of meningioma, is distinguished by losses in Chr 1p and 22q^8,16,23^. In contrast, MenG B meningiomas typically display Chr 22q deletions^8,16,23^. Given that bulkRF-based classifiers have previously shown significant confusion in distinguishing patient samples between MenG C and A tumors, large-scale copy number variation (CNV) signals are crucial in the accurate classification of meningioma classes. In previous work, we demonstrated how CNV profiles can be inferred from bulk transcriptomics and the utility of these profiles in the accurate classification of tumors^24^. Leveraging our previously published tool, CaSpER^24^, we generated RNA-inferred CNV profiles for each sample and employed them in training a CNV-based Random Forest classifier (cnvRF).

Finally, the outputs of the bulkRF, ctRFs, and cnvRF, which are probabilities that a given sample of each possible class, were used as features in a Random Forest model. Thus, the scRFs, cnvRF, and bulkRF are integrated into a meta-model that is used to assign a sample’s final classification.

##### Classifiers trained with features selected from bulk transcriptomics perform well but confuse benign and malignant samples

We first performed pairwise differential expression analysis between the meningioma groups within the training cohort. A well-labeled training cohort is essential for developing accurate, reliable, and interpretable machine-learning models for classification tasks. Therefore, building upon our previous work^4^, we identified the patient samples that were consistently classified using multi-omics profiling. These samples, prototypical members of their meningioma groups, were established as the training cohort for the study. Differential analysis was employed to identify class-specific differentially expressed genes (DEGs). Using these group-specific DEGs, we trained a Random Forest classifier (RF parameters).

We next extracted the samples (n=7) that exhibited consistent classification across other multi-omics datasets but did not align with the bulk transcriptomics class initially identified in our previous study^4,9^. Our updated bulk model (bulkRF), trained with well-characterized samples, outperformed the initial model presented in Patel et al.’s study. However, upon closer examination, we observed that the bulk-RF model faced challenges in reliably distinguishing between MenG A and C tumors (**Figure 2 a-c**). The samples that show confusion between groups A and C are typically found near the boundary between the A and C clusters in the PCA and tSNE plot (**Figure 2a-b, Supp Figure 1**). Moreover, some of the misclassified A samples have a higher CNV burden (**Supp Figure 2**).

**Figure 2.**
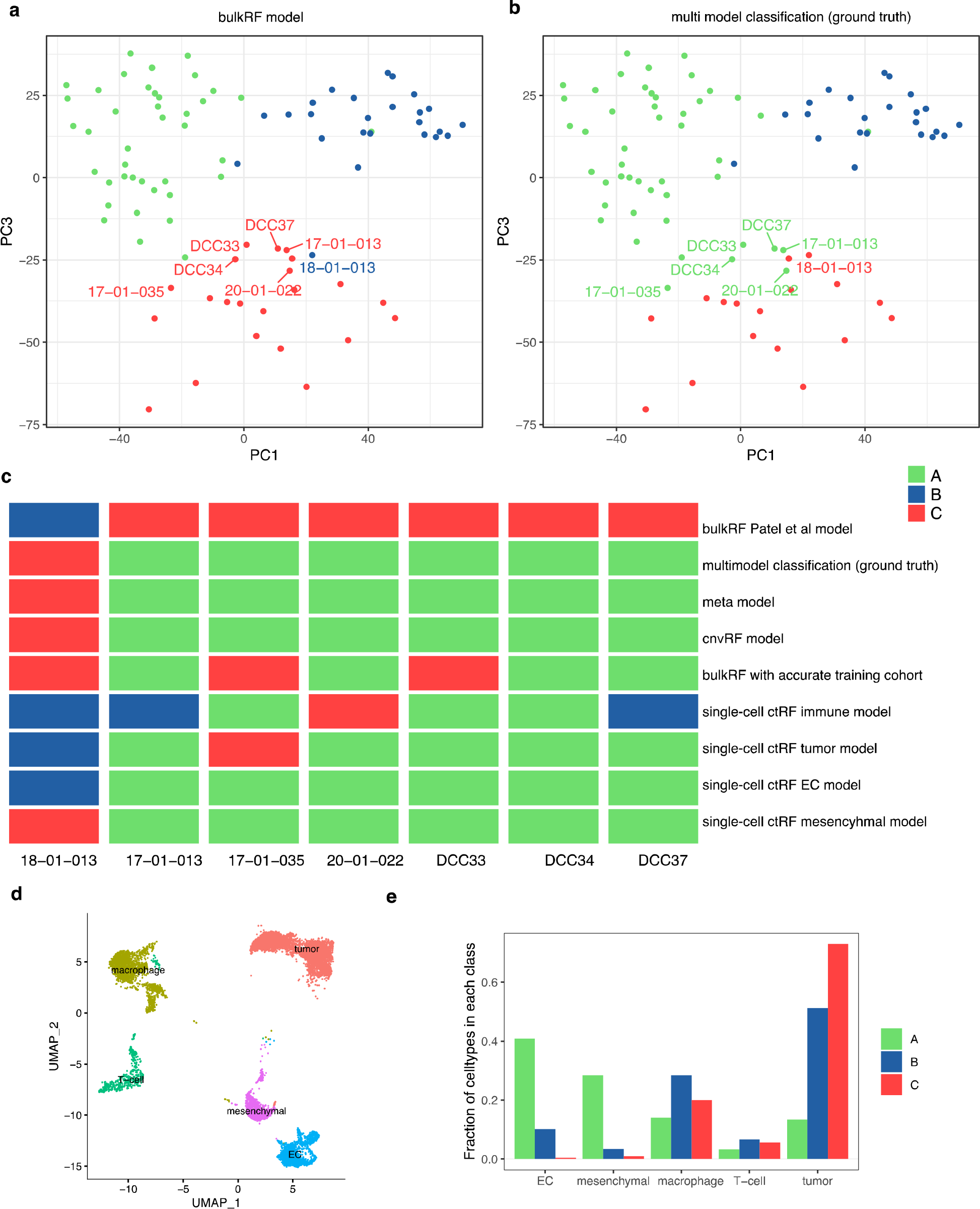
PCA Plot of in-house meningioma bulk data. Each point’s color indicates the class prediction: (a) by the bulkRF model, and (b) by the meta-model. Samples with consistent classification across multi-omics datasets but differing from the initial bulk transcriptomics class determined in our prior study by Patel et al., are labeled. (c) Oncoprint detailing the predictions of each sample from each model is shown. (d) Clustering of single cells with corresponding annotated cell types is shown. (e) Proportion of each cell type within individual classes is shown.

##### Single-cell models and CNV models resolve benign-malignant confusion

Single-cell sequencing offers unparalleled resolution in the molecular characterization of tumor samples^25^. Interestingly, tumor samples previously classified as group A were enriched in endothelial and mesenchymal cells, those classified as B were enriched in macrophages, and tumors of both B and C were enriched in tumor cells (**Figure 2d-e, Supp Figure 3-6**).

To explore the potential of this enhanced resolution in distinguishing between different molecular groups of meningioma, we trained a classifier using features extracted from single-cell data. We used single-cell sequencing data from n=6 meningioma samples (~10K cells) with annotated tumor classifications and cell types for each sequenced cell^10,26^ (**Supp Figure 3-6)**. Then, we identified group-specific markers for each cell type by performing DEG analysis using Seurat R package. Using the expression levels of these markers in bulk transcriptomic data, we trained cell-type specific Random Forest models.

We again extracted a subset of samples (n=7) that displayed concordant classification across other multi-omics datasets but differed from the initial bulk transcriptomics class identified in our prior study^4,9^ (**Figure 2c**). Among the models developed, those based on models generated from the mesenchymal cell (mesenchyme-RF) and EC cell (EC-RF) markers exhibited superior performance, demonstrating reduced misclassification between MenG A and C samples (**Figure 2c**).

These results suggest that single-cell models capture features that are important for differentiating between MenG A and C meningioma that are not evident in the bulk data, but that these features, alone, are not sufficient to generate models that can outperform the bulk model. Similarly, the cnvRF model, which relies on RNA-inferred CNV signals, performed well in resolving the confusion between benign and malignant classifications (**Figure 2c**).

### Integrating bulk, CNV, and single-cell-based classifiers yields optimal performance and provides superior clinical prognoses

To determine whether single-cell models and the bulk-RF could be synergistically integrated, we stacked 6 models (mesenchymal ctRF, immune ctRF, tumor ctRF, EC ctRF, bulkRF, cnvRF) into a meta-model. The meta-model was constructed by using the predictions of the component models as a feature for a Random Forest that generated the final predictions for each sample.

Next, we evaluated the performance of our CLIPPR algorithm meta-model on a bulk meningioma dataset consisting of 792 samples gathered from multiple institutions^10,27–29^. We performed class predictions on this integrated data (n=792) using the models trained on the well-characterized training cohort^4^. We assessed the performance of CLIPPR by concentrating on the samples that exhibited inconsistencies between the bulkRF model and the CLIPPR model (meta-model) (**Figure 3a-c, Supp Figure 7**). Given that the ground truth classes are unavailable for all 792 samples, we assessed the performance of CLIPPR by inspecting the Kaplan-Meier curves of the samples that exhibited inconsistencies between the bulkRF model and the CLIPPR model. Kaplan-Meier analysis reveals significant confusion between benign and malignant samples within the inconsistently classified samples by both the bulkRF and CLIPPR models, as defined by their recurrence rates. Among samples exhibiting inconsistencies between the bulkRF model and the CLIPPR meta-model (n=53), those classified as MenG A in the bulkRF model displayed higher recurrence rates than those classified as MenG C in the bulkRF model (p < .008). In contrast, within the subset of samples displaying inconsistencies between the bulkRF model and the CLIPPR meta-model, the classifications from the CLIPPR meta-model algorithm provided classifications concordant with tumor behavior. Specifically, samples displaying inconsistencies categorized as MenG C in the CLIPPR meta-model demonstrated notably higher rates of recurrence compared to those classified as MenG A (p < .02) (**Figure 3d-e**). With the end goal of using this meningioma classification scheme in the clinic, it was important to assess whether improved discrimination between MenG A and C tumors translated to improved clinical prognostication.

**Figure 3.**
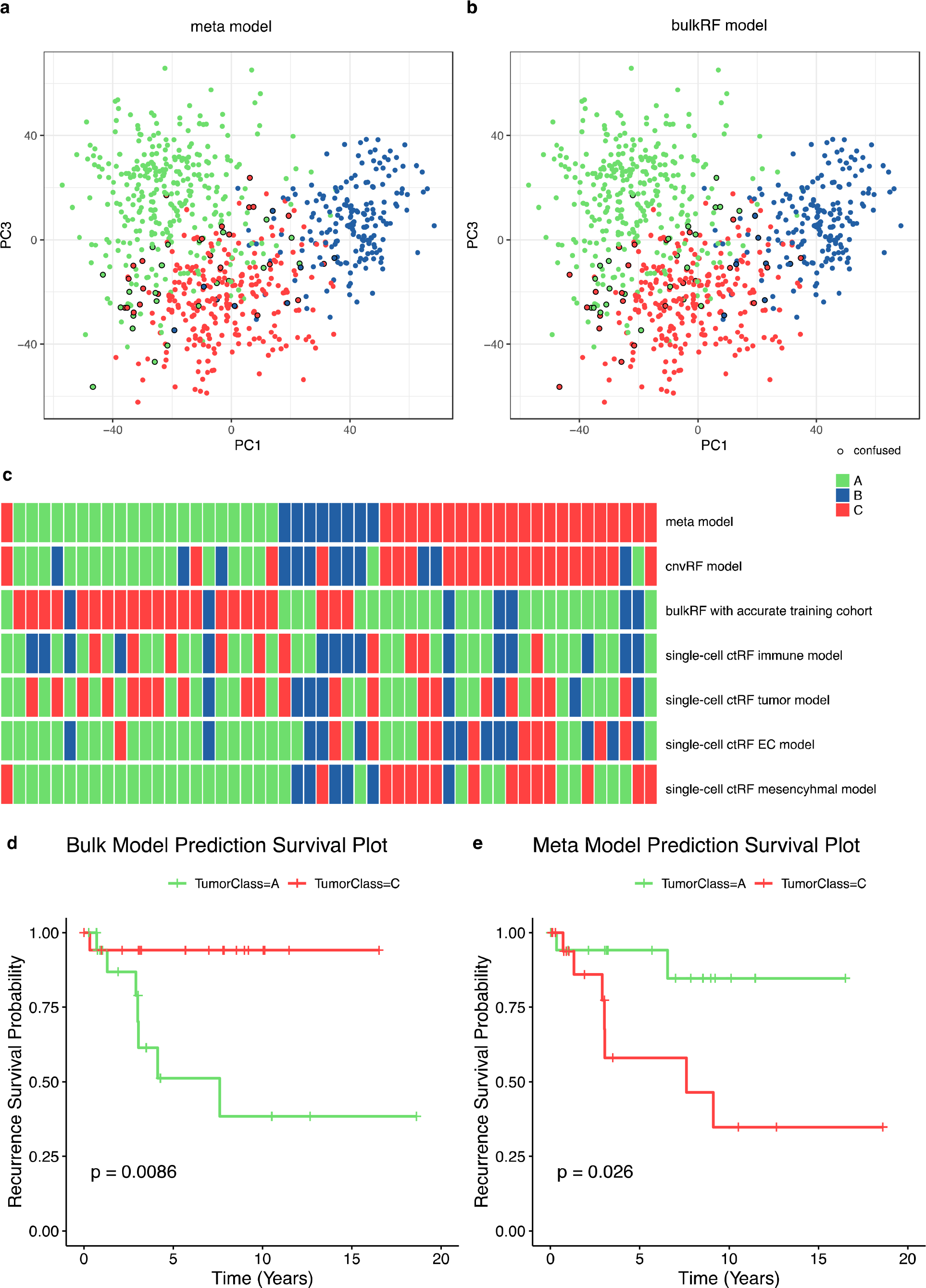
PCA Plot of meningioma bulk data from various institutions (n=792). Each point’s color represents sample class prediction: (a) by the CLIPPR meta model, and (b) by the bulkRF model. Samples with classification discrepancies between the two models are circled in black (n=53). (c) Oncoprint illustrating the predictions for each inconsistently classified sample by both models (n=53). (d) Kaplan Meier plot based on bulkRF model predictions, highlighting poorer survival of benign (type A) samples relative to malignant (type C) samples (p-value: 0.008). (e) Kaplan Meier plot using CLIPPR meta model predictions, showing expected poorer survival for malignant (type C) samples compared to benign (type A) samples (p-value: 0.02).

#### CLIPPR algorithm demonstrates generalizability with application to GBM

We aimed to demonstrate the generalizability of the CLIPPR algorithm in the context of glioma data. Specifically, we utilized subtype-specific cell-type features extracted from our in-house single-cell glioma dataset, which includes patients with IDH Mutant (astrocytoma), IDH Mutant (oligodendroglioma), and IDH Wild-Type (WT) tumor classes^22^ (**Figure 4a**). Subsequently, we partitioned the bulk RNA-Seq data from TCGA into both validation and training cohorts. We constructed our models using features extracted from CNV, bulk transcriptomics, and single-cell type-specific markers. In our analysis, we assessed the prediction accuracy of the sample classes in the validation cohort by comparing the bulkRF and meta-model predictions. Our analysis showed that the meta-model exhibited superior concordance than the bulkRF model (**Figure 4b)**.

**Figure 4.**
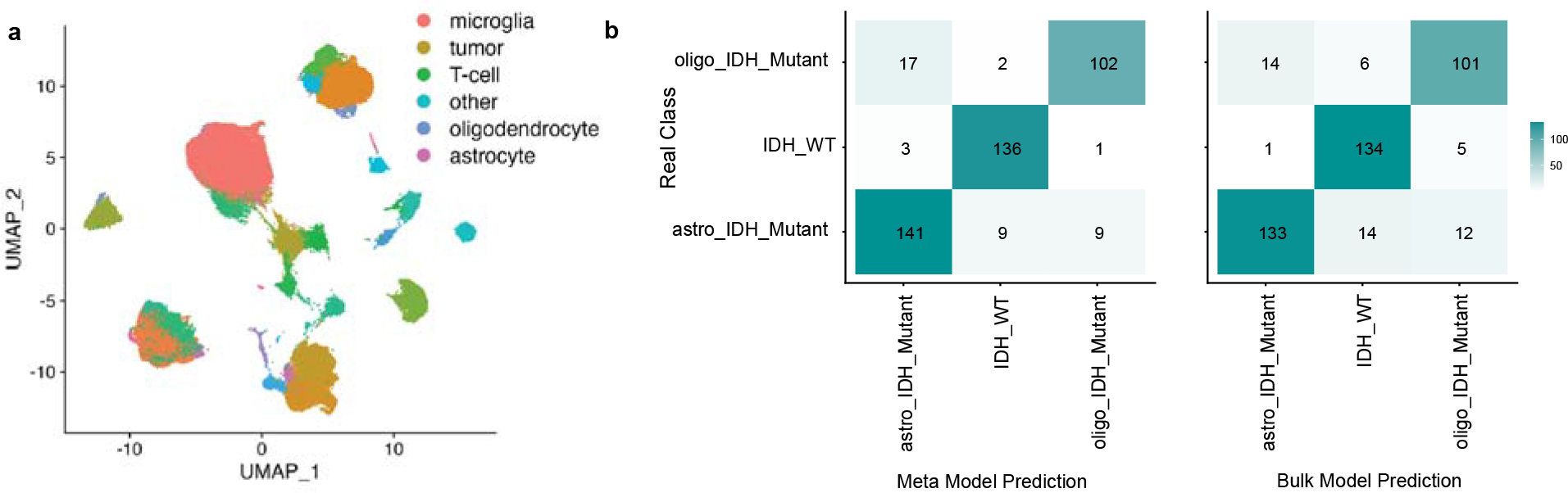
(a) Clustering of glioma single cells, with each cluster annotated by cell type. (b) Comparative prediction accuracy of sample classes between the CLIPPR meta model and the bulkRF model, with the CLIPPR meta-model exhibiting better accuracy.

## Discussion

Previously, we adopted an integrative multi-omics profiling strategy to molecularly classify meningioma^4^. This approach led to the discovery of three distinct molecular groups (A, B, and C) that surpassed the traditional WHO grading system in predicting recurrence risks^4^. Half of MenG C tumors will recur after only 47 months, despite the fact that the majority in our cohort are WHO grade I tumors^30^. However, classifiers relying solely on bulk transcriptomic data have shown significant confusion in distinguishing MenG A from MenG C tumors. Here we developed CLIPPR, a method that predicts the meningioma classes by leveraging single-cell data and RNA-inferred CNV signal to enhance the prediction accuracy of bulk data classifiers. We demonstrate that using models trained on features learned from single-cell data accurately resolved the confusion between MenG A and C tumors but had limited overall accuracy.

Similarly, models generated from RNA-inferred large-scale CNV signals also predicted malignant class accurately with limited overall accuracy. However, combining the top-performing single-cell models, CNV models with the initial bulk model into a meta-model resulted in the strongest performance, with superior overall accuracy and better benign-malignant resolution.

In summary, here we showed that CLIPPR distinguishes between clinically benign and malignant molecular classes more accurately in meningiomas. Our method can be easily adapted for application to different cancer types. Moreover, our meningioma and glioma models can be downloaded and used in other facilities to predict the tumor class of new patient samples for clinical use or to analyze in-house bulk RNA-Seq data.

## Methods

### Single-cell processing

We ran samples on the 10X Chromium platform to produce next-generation sequencing libraries. We performed standard procedures for filtering, mitochondrial gene removal, and variable gene selection using the Seurat pipeline^31^. The criteria for cell/gene inclusion were as follows: genes present in more than three cells were included, cells that expressed >300 genes were included, the number of genes detected in each cell was >200 and <5000, and the mitochondria ratio was 10. We integrated cells from different patients using the Harmony algorithm^32^. Next, we visualized clusters using a uniform manifold approximation and projection constructed from the Harmony-corrected PCA. This visualization was created using the *runUMAP, FindNeighbors*, and *FindClusters* functions of the Seurat package. We extracted differentially expressed genes among clusters using *FindAllMarkers* function of Seurat package^31^. Next, we employed well-established cell type markers to annotate each cluster with its corresponding cell type. Additionally, we integrated our CNV-calling algorithm, CaSpER^24^, to precisely identify tumor cells at a single-cell resolution. To identify large-scale CNV events, which we defined as involving at least one-third of a chromosomal arm. Visualizations with UMAP plots were employed to validate annotations.

### Analyzing meningioma and glioma bulk expression data and survival analysis

We processed raw reads from the in-house meningioma dataset, as well as RNA-seq data from two different institutes, using a custom pipeline that incorporates FastQC and RSeQC for read and alignment quality assessment. Reads were aligned to the GRCh38 Human reference genome, followed by mapping to the human transcriptome based on UCSC gene annotations using STAR tool^33^. Next, an expression count matrix was generated through our in-house pipeline. The RNA-seq read counts for genes were then normalized, and applied a variance stabilizing transformation, and differential gene expression analysis was performed using the DESeq2 package, with correction for institute-specific effects.

TCGA-GBM (high grade glioma), TCGA-LGG (low grade glioma) raw read counts and accompanying clinical data are downloaded using TCGAbiolinks R package^34^. TCGA-GBM, TCGA-LGG and our bulk RNA-Seq data of the IDH Mutant cohort were both normalized and variance stabilizing transformation was applied using the DESeq2 package^35^.

The predicted groups are compared against recurrence in a Cox Proportional Hazards (Cox) survival model. We used *survminer* and *survival* R package for the survival analysis.

### Feature Selection

Feature selection is a critical component in the construction of classification. The CLIPPR algorithm, which is comprised of three parallelized classifiers, employs three feature selection schemes that each correspond to the data respective data of the component classifiers: bulk, single-cell and CNV.

The bulk sequencing feature selection scheme utilizes pairwise differential expression between the transcriptomes of the meningioma sub-groups. Differentially expressed genes (DEGs) that were statistically significant (p adj. < .05), had a log2fold change with an absolute value greater than 1.5 and were specific to pairwise comparisons for one sub-group, were utilized as features for the bulk model.

The single-cell feature selection scheme employs a similar approach to identifying class-specific features. First, sequencing data is stratified by cell-type, then by class. Next the FindMarkers function from the Seurat package is used to identify markers that correspond to a specific cell-type within a meningioma sub-group.

We used the CaSpER^24^ algorithm for CNV feature selection in our study. Specifically, we used CaSpER to perform signal smoothing on the expression count matrix. Following the smoothing process, we obtained the smoothed expression signals and subsequently computed the median signal for each chromosome arm, focusing on signals from 1p, 14q, and 22q of each sample. We chose to focus on 1p, 14q, and 22q because they represent the most prevalent CNV events within MenG C, while 22q is the only chromosomal deletion event in MenG B.

### Model Generation

The Random Forest models utilized in the CLIPPR algorithm were created using the R package randomForest with the ntree parameter 5000 using the randomForest function. We constructed separate random forest models using the features described above, which were selected from CNV signals, DEGs in bulk, and DEGs specific to cell type classes in single-cell data. Next, we used metamodels to harness the power of ensemble learning by utilizing the predictions generated from individual models. The metamodel combine the predictions of each model to create a comprehensive final model. This ensemble approach leverages the strengths of each individual model, improving the overall predictive performance and robustness of our analysis, ultimately enhancing our ability to make accurate and reliable inferences in our research.

## Data availability

The accession numbers for the previously published bulk and single-cell meningioma RNA-seq data used in the study are GSE221536, GSE213544, GSE212666 and GSE183653^10,22,26,29^.

## Authors contribution

A.S.H., A.J.P., A.S., and A.O.H. are responsible for the conception of this project, the study and pipeline design, and the interpretation of the results. A.S.H., A.S. wrote the code for the CLIPPR pipeline and performed the analysis. S.W. performed the analysis. A.S.H., A.S., and A.O.H. prepared the manuscript. A.J.P., T.J.K, A.B.K, C.W.E., S.H.N., S.T.M, D.R.R, and S.W. contributed to the manuscript with feedback from all authors.

## References

1. DeAngelis, L. M. Brain tumors. N. Engl. J. Med. 344, 114–123 (2001).

2. Prados, M. D. et al. Toward precision medicine in glioblastoma: the promise and the challenges. Neuro. Oncol. 17, 1051–1063 (2015).

3. Louis, D. N. et al. The 2021 WHO classification of tumors of the Central Nervous System: A summary. Neuro. Oncol. 23, 1231–1251 (2021).

4. Bayley, J. C., 5th. et al. Multiple approaches converge on three biological subtypes of meningioma and extract new insights from published studies. Sci. Adv. 8, eabm6247 (2022).

5. Ostrom, Q. T. et al. CBTRUS statistical report: Primary brain and other central nervous system tumors diagnosed in the United States in 2016-2020. Neuro. Oncol. 25, iv1–iv99 (2023).

6. Wiemels, J., Wrensch, M. & Claus, E. B. Epidemiology and etiology of meningioma. J. Neurooncol. 99, 307–314 (2010).

7. Rogers, L. et al. Meningiomas: knowledge base, treatment outcomes, and uncertainties. A RANO review. J. Neurosurg. 122, 4–23 (2015).

8. Harmanci, A. S. et al. Integrated genomic analyses of de novo pathways underlying atypical meningiomas. Nat. Commun. 8, 14433 (2017).

9. Patel, A. J. et al. Molecular profiling predicts meningioma recurrence and reveals loss of DREAM complex repression in aggressive tumors. Proc. Natl. Acad. Sci. U. S. A. 116, 21715–21726 (2019).

10. Choudhury, A. et al. Meningioma DNA methylation groups identify biological drivers and therapeutic vulnerabilities. Nat. Genet. 54, 649–659 (2022).

11. Choudhury, A. et al. NOTCH3 drives meningioma tumorigenesis and resistance to radiotherapy. bioRxiv 2023.07.10.548456 (2023) doi:10.1101/2023.07.10.548456.

12. Harmanci, A. S. et al. Aggressive human MenG C meningiomas have a molecular counterpart in canines. Acta Neuropathol. 147, 42 (2024).

13. Zakimi, N. et al. Canine meningiomas are comprised of 3 DNA methylation groups that resemble the molecular characteristics of human meningiomas. Acta Neuropathol. 147, 43 (2024).

14. Olar, A. et al. Global epigenetic profiling identifies methylation subgroups associated with recurrence-free survival in meningioma. Acta Neuropathol. 133, 431–444 (2017).

15. Sahm, F. et al. DNA methylation-based classification and grading system for meningioma: a multicentre, retrospective analysis. Lancet Oncol. 18, 682–694 (2017).

16. Clark, V. E. et al. Genomic analysis of non-NF2 meningiomas reveals mutations in TRAF7, KLF4, AKT1, and SMO. Science 339, 1077–1080 (2013).

17. Clark, V. E. et al. Recurrent somatic mutations in POLR2A define a distinct subset of meningiomas. Nat. Genet. 48, 1253–1259 (2016).

18. Brastianos, P. K. et al. Genomic sequencing of meningiomas identifies oncogenic SMO and AKT1 mutations. Nat. Genet. 45, 285–289 (2013).

19. Bi, W. L. et al. Genomic landscape of high-grade meningiomas. NPJ Genom. Med. 2, (2017).

20. Raleigh, D. et al. Targeted gene expression profiling predicts meningioma outcomes and radiotherapy responses. Res. Sq. (2023) doi:10.21203/rs.3.rs-2663611/v1.

21. Ballester, L. Y., Olar, A. & Roy-Chowdhuri, S. Next-generation sequencing of central nervous systems tumors: the future of personalized patient management. Neuro-oncology vol. 18 308–310 (2016).

22. Curry, R. N. et al. Glioma epileptiform activity and progression are driven by IGSF3-mediated potassium dysregulation. Neuron 111, 682-695.e9 (2023).

23. Patel, A. J. et al. Molecular profiling predicts meningioma recurrence and reveals loss of DREAM complex repression in aggressive tumors. bioRxiv (2019) doi:10.1101/679480.

24. Harmanci, A. S., Harmanci, A. O. & Zhou, X. CaSpER identifies and visualizes CNV events by integrative analysis of single-cell or bulk RNA-sequencing data. Nature Communications vol. 11 Preprint at 10.1038/s41467-019-13779-x (2020).

25. Zhang, Y. et al. Single-cell RNA sequencing in cancer research. J. Exp. Clin. Cancer Res. 40, (2021).

26. Harmanci, A., Harmanci, A. S., Klisch, T. J. & Patel, A. J. XCVATR: detection and characterization of variant impact on the Embeddings of single-cell and bulk RNA-sequencing samples. BMC Genomics 23, 841 (2022).

27. Vasudevan, H. N. et al. Intratumor and informatic heterogeneity influence meningioma molecular classification. Acta Neuropathol. 144, 579–583 (2022).

28. Nguyen, M. P. et al. Supervised machine learning algorithms demonstrate proliferation index correlates with long-term recurrence after complete resection of WHO grade I meningioma. J. Neurosurg. 138, 86–94 (2023).

29. Choudhury, A. et al. Hypermitotic meningiomas harbor DNA methylation subgroups with distinct biological and clinical features. Neuro. Oncol. 25, 520–530 (2023).

30. Khan, A. B. et al. Even heterozygous loss of CDKN2A/B greatly accelerates recurrence in aggressive meningioma. Acta Neuropathol. 145, 501–503 (2023).

31. Stuart, T. et al. Comprehensive integration of single-cell data. Cell 177, 1888-1902.e21 (2019).

32. Korsunsky, I. et al. Fast, sensitive and accurate integration of single-cell data with Harmony. Nat. Methods 16, 1289–1296 (2019).

33. Dobin, A. et al. STAR: ultrafast universal RNA-seq aligner. Bioinformatics 29, 15–21 (2013).

34. Mounir, M. et al. Analyses of cancer data in the Genomic Data Commons Data Portal with new functionalities in the TCGAbiolinks R/Bioconductor package. bioRxiv (2018) doi:10.1101/350439.

35. Love, M. I., Huber, W. & Anders, S. Moderated estimation of fold change and dispersion for RNA-seq data with DESeq2. Genome Biol. 15, 550 (2014).

